# TIVAN-indel: A computational framework for annotating and predicting noncoding regulatory small insertion and deletion

**DOI:** 10.1101/2022.09.28.509993

**Authors:** Aman Agarwal, Li Chen

## Abstract

**Motivation:** Small insertion and deletion (sindel) of human genome has an important implication for human disease. One important mechanism for noncoding sindel to have an impact on human diseases and phenotypes is through the regulation of gene expression. Nevertheless, current sequencing technology may lack statistical power and resolution to pinpoint the causal sindel due to lower minor allele frequency or small effect. As an alternative solution, a supervised machine learning method can identify the otherwise missing causal sindels by predicting the regulatory potential of sindels directly. However, computational methods for annotating and predicting the regulatory sindels, especially in the noncoding regions, are underdeveloped.

**Results:** By leveraging recognized sindels in *cis*-expression quantitative trait loci (*cis*-eQTLs) across 44 tissues and cell types in GTEx, and a compilation of both generic functional annotations and tissue/cell typespecific multi-omics features generated by a sequence-based deep learning model, we developed TIVAN-indel, which is an XGBoost-based supervised framework for scoring noncoding sindels based their potential to regulate the nearby gene expression. As a result, we demonstrate that TIVAN-indel achieves the best prediction performance in both cross-validation with-tissue prediction and independent cross-tissue evaluation. As an independent evaluation, we train TIVAN-indel from “Whole Blood” tissue in GTEx data and test the model using 15 immune cell types from an independent study DICE. Lastly, we perform an enrichment analysis for both recognized and predicted sindels in key regulatory regions such as chromatin interactions, open chromatin and histone modification sites, and find biologically meaningful enrichment patterns.

**Availability and implementation:** https://github.com/lichen-lab/TIVAN-indel

## 1 Introduction

Only after single nucleotide variants (SNVs), small insertion and deletion (sindels) (e.g., <50bp) are the second most common mutations in the human genome, which make up 15–21% of human polymorphism [1]. In the coding regions, sindels can alter the protein sequence by adding/repathogenicmoving multiples of three nucleotides, which leads to the insertion/deletion of one or more amino acids (non-frameshift sindels) or adding/removing a number of base pairs that is not a multiple of three, which consequently disrupts the triplet reading frame of a DNA sequence (frameshift sindels). Recent studies have shown that coding sindels can significantly influence a range of phenotypic and molecular effects [2, 3]. For example, cystic fibrosis, one common human disease, is frequently caused by a coding sindel within the CFTR gene that eliminates a single amino acid [4]. Moreover, noncoding sindels (nc-sindels) have also been found to play an important role in human traits and diseases by affecting the gene expression such as alter the pattern, phasing and spacing of DNA sequences within promoters [5]. One example is that DNA insertions within the promoter region of the FMR1 gene cause Fragile X syndrome in humans [6]. However, the functional consequence of nc-sindels are less explored compared to their coding counterparts.

Despite the experimental validation and clinical interpretation for sindels in the past decades, the knowledge of sindels is mostly limited to thousands of pathogenic sindels in HGMD [7] and ClinVar [8]. However, considering a total of discovered 1.2 million indels from 125,748 exomes, and 33 million indels from 15,708 genomes in gnomAD v2.1 [9], the functional consequences of most indels are still unknown. To achieve a genome-wide prediction for the functional consequence of sindels in the noncoding regions, several computational tools have been developed. CADD adopts SVM to integrate diverse genomic annotations such as sequence context, evolutionary constraints, epigenetic measurements for observed and simulated sindels [10] into a single score, which ranks the deleteriousness of sindels in both coding and noncoding regions. FATHMM-indel, another SVM-based machine learning approach, adopts a similar training feature set as CADD, to exclusively predict pathogenic sindels in the noncoding regions [11] by using pathogenic noncoding sindels from HGMD as training samples. Recently, deep learning methods have been developed to predict functional noncoding variants, which include both SNV and sindel. The representative work such as DeepSEA [12] and DanQ [13] can use genomic sequence to simultaneously predict thousands of chromatin-profiling data, which include transcription factor binding, histone-mark profiles and open chromatin across multiple cell types. Using the predicted chromatin-profiling data under between reference and alternative alleles as genomic annotations, DeepSEA adopts a boosted logistic regression classifier to discriminate noncoding pathogenic sindels in HGMD and background sindels from 1000 Genomes [14].

Regulation of gene expression is a key molecular mechanism for noncoding sindels to achieve phenotypic and clinical significance by disrupting the normal sequence pattern in the promoters and enhancers, which can influence the nearby gene expression [6, 5, 15]. Similar to SNVs, the same challenge is faced by current sequencing experiments to identify nc-indels that regulate gene expression due to the small sample size, low minor allele frequencies (MAFs), small effects or missing gene expression data for matched genotype data. Moreover, existing computational methods, such as CADD, FATHMM-indel and DeepSEA focus on predicting the pathogenicity or deleteriousness of nc-sindels, rather than the molecular impact. Therefore, it is crucial to develop a computational tool for predicting the functional potential of nc-sindels that may alter gene expression, which can provide a mechanistic understanding of the functional role of nc-indels in disease pathogenesis.

In this work, we develop a novel computational framework, named TIVAN-indel (<u>TI</u>ssue-specific <u>V</u>ariant <u>A</u>nnotation for <u>N</u>oncoding <u>indel</u>), which aims to predict nc-sindels that regulate gene expression. Our contribution mainly lies on two aspects. First, to our best knowledge, this is the first computational tool to predict tissue/cell type-specific regulatory nc-sindels. Second, TIVAN-indel leverages both generic CADD annotations (e.g., evolutionary score, sequence characteristics) and large-scale tissue/cell type-specific multi-omics features derived from a sequence-based deep learning model (i.e., DanQ) as the functional annotations for nc-sindels in a supervised machine learning framework to improve the prediction performance. We benchmark TIVAN-indel and existing approaches on 44 tissues/cell types in GTEx using a cross-validation within-tissue approach and an independent cross-tissue approach respectively. As an independent evaluation, we train TIVAN-indel from “Whole Blood” tissue in GTEx data and test the model using 15 immune cell types from an independent study DICE. To demonstrate nc-sindels play a key role in gene regulation, we perform an enrichment analysis for both labeled and predicted nc-sindels in key regulatory regions such as chromatin interactions, open chromatin and histone modification sites, and find biologically meaningful enrichment patterns.

## 2 Methods and Materials

### 2.1 Creating the training samples

Using a similar approach as previously developed TIVAN [16], we obtain tissue/cell type-specific nc-sindels in *cis*-QTLs that influence nearby gene expression and create positive and negative nc-sindels from Genotype-Tissue Expression (GTEx) [17]. For each tissue/cell type, to create the positive set, we select nc-sindels from GTEx (v6p) that meet three criteria including (i) the length of nc-sindels less than 100bp; (ii) distance to transcription start site (TSS) less than 100kb as a nc-sindel may have an impact on a nearby gene; (iii) q-value less than 0.05 with at least one associated gene. To create the negative set, we obtain all nc-sindels from GTEx (v7) and perform a series of filtering criteria, which include (i) non-overlap with any nc-sindels in the positive set; (ii) associated with at least one gene in the positive set; (iii) distance to the TSS less than 100kb; (iv) q-value larger than 0.2; (v) with a matched empirical distribution of minor allele frequency (MAF) as nc-sindels in the positive set. Without loss of generality, we create a balanced negative set. The sample sizes of the positive sets across 44 tissues/cell types are summarized in Table S1.

### 2.2 Creating the functional annotations

We create functional data from two sources for annotating nc-sindels. The first source is the precomputed functional annotations for 48 million indels from the latest CADD v1.6 (Genome build GRCh37/hg19) [18]. We only keep CADD annotations with less than 10% missing values and impute the missing values using the median of non-missing values for each annotation. As a result, we have 45 functional annotations, which mainly include (i) base-level features such as conservation score and GC content; (ii) chromatin states in 48 cell types; (iii) H3K27ac, H3K4me1 and H3K4me3 levels; (iv) number of frequent, rare and single occurrence TOPMed SNVs in BRAVO server [19] at 100, 1000 and 10000 bp resolution. The detailed descriptions of the 45 functional annotations can be found in Table S2.

The second source is the tissue/cell type-specific multi-omics features, which are derived from the sequence-based deep learning model. First, we obtain a fixed length of 1000bp reference genomic sequence of each sindel. Similarly, we create the alternative sequence by changing the reference allele to the alternative allele. Each genomic sequence is one-hot encoded as ‘A’ - [1, 0, 0, 0], ‘G’ - [0, 1, 0, 0], ‘C’ - [0, 0, 1, 0], and ‘T’ - [0, 0, 0, 1]. Next, we exploit DanQ [13], a recently developed multi-task convolutional neural network, which takes the one-hot encoded reference/alternative genomic sequence as the input to simultaneously predict large-scale chromatin-profiling data, including transcription factor binding, histone-mark profiles and open chromatin across multiple cell types for each nc-sindel. As a result, DanQ outputs 919 binary chromatin profiles, which are derived from the ENCODE and Roadmap Epigenomics, including TF binding site (690), DNase I hypersensitive site (125) and Histone modification sites (104) [13]. A total of 919+919=1838 predicted chromatin profiles can be used to annotate for reference and alternative alleles of the sindel respectively. It should be noted that other deep learning models such as DeepSEA [12] can also be used for the same purpose. We adopt DanQ in this work as it has been demonstrated to outperform DeepSEA in the prediction task [13]. Collectively, both 45 generic functional annotations and 1838 multi-omics features are used to annotate the labeled nc-sindels and train TIVAN-indel.

### 2.3 Computational framework of TIVAN-indel

As shown in (Figure 1), the computational framework of TIVAN-indel consists of two steps. The first step is feature extraction. Given the genomic loci of a nc-sindel, TIVAN-indel will obtain 45 precomputed CADD functional annotations and 919 multi-omics features predicted by DanQ under reference and alternative sequence respectively. The second step is model training. Once nc-sindels are annotated with the extracted features, TIVAN-indel will adopt XGBoost as the supervised approach to train the prediction model. It should be noted that TIVAN-indel is trained in a tissue/cell type-specific way as nc-sindels influence gene expression differently in different tissue/cell types.

**Figure 1:**
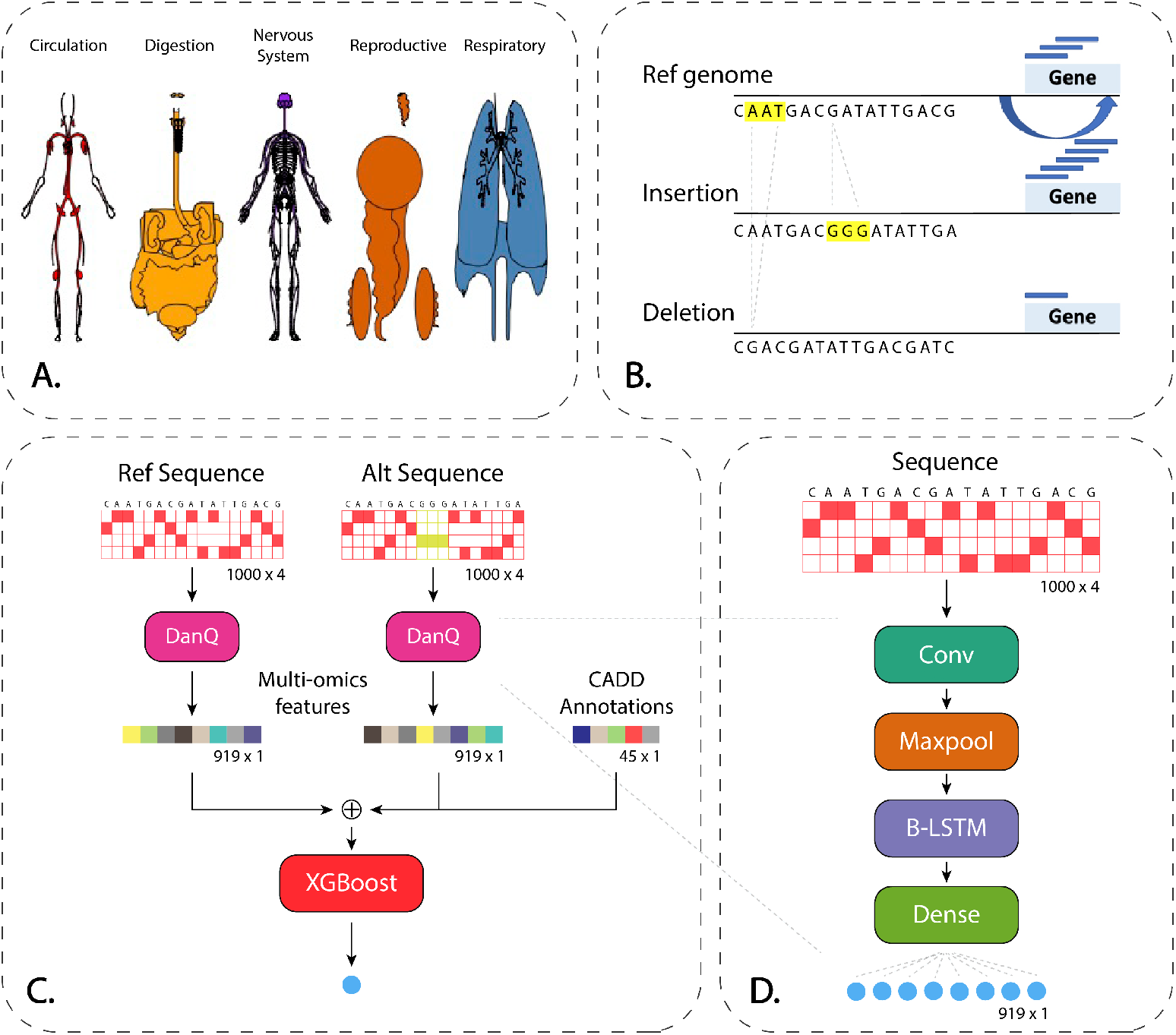
Overview of TIVAN-indel. **A.** We collect nc-sindels in *cis*-eQTLs from 44 tissues/cell types from GTEx. **B.** A cartoon example to show a small insertion/deletion in *cis*-eQTLs can affect gene expression. **C.** TIVAN-indel takes both precomputed CADD annotations and predicted multi-omics features under reference and alternative genomic sequences by DanQ to annotate labeled nc-sindels. TIVAN-indel adopts XGBoost for the binary classification. **D.** A brief illustration of the model architecture of DanQ.

### 2.4 Promoter-centered captured HiC data

To calculate the enrichment of nc-sindels in tissue-matched chromatin interactions, we collect and curate a deposition of promoter-centered chromatin interactions in 3D-genome Interaction Viewer and database (3DIV) [20], which has profiled a comprehensive set of promoter-centered capture Hi-C datasets (pcHi-C) across 27 human tissue/cell types in 4 categories including embryonic stem cell, 4 early embryonic lineages, 2 primary cell lines and 20 primary tissue types. For each tissue/cell type, pcHi-C provides both promoter-enhancer interactions (PE) and promoter-promoter interactions (PP). Next, we adopt a similar procedure to create positive and negative chromatin interactions as our previous work [21]. Specifically, we filter chromatin interactions with distance more than 10^6^ and consider only intra-chromosomal chromatin interactions. We further format each anchor into 2kb region with 1kb upstream and downstream of the midpoint of the anchor. Among all interactions, we label significant interactions (FDR<0.1) as the positive set and only keep tissue/cell type with more than 100 significant interactions. As a result, the number of positive samples ranges from 972 to 56057 with a median of 5071 among 26 tissues/cell types for PE and from 187 to 10338 with a median of 917 among 25 tissues/cell types for PP. To create the negative set, we select interactions (FDR>0.5) with matched GC contents as the positive set. Without loss of generality, the number of negative interactions is set as the same as the positive interactions. More details of matched tissues/cell types of GTEx and tissues/cell types of pcHi-C in 3DIV can be found in Table S3.

### 2.5 Roadmap Epigenomics data

To calculate the enrichment of nc-sindels in tissue-matched epigenomic regions, we collect tissue/cell type-specific epigenomic datasets from Roadmap Epigenomics [22]. Especially, we collect ChIP-seq data related to open chromatin and key histone marks. For active histone marks, we collect H3K4me3 and H3K9ac associated with promoter activation, H3K4me1 and H3K27ac associated with enhancer activation, H3K36me3 associated with active gene body [23] and H3K27me3 and H3K9me3 associated with gene repression. To create the positive set, we keep significant ChIP-seq peak regions (FDR<0.05) as the positive set and fix the peak size into 1000bp window by extending the peak center upstream and downstream 500bp. To create the negative set, we random choose the same number of 1000bp windows across the whole genome with a matched GC content distribution as the positive set. More details of matched tissues/cell types of GTEx and tissues/cell types of ChIP-seq data from chosen histone marks and open chromatin in Roadmap Epigenomics can be found in Table S4.

## 3 Results

### 3.1 Integrating both CADD functional annotations and multi-omics features derived from sequence-based deep learning model improves the prediction for regulatory noncoding small indels

Conventional computational methods for predicting regulatory nc-sindels such as CADD and FATHMM-indel mainly adopt tens of genomic annotations as the training features. With the advent and availability of large-scale tissue/cell type-specific epigenomic data deposited in multiple consortia such as ENCODE [24] and Roadmap Epigenomics as well as the rationale that regulatory nc-sindels can perform a function in the epigenomic regions, we hypothesize that curating these multi-omics data into additional training features can improve the prediction for regulatory nc-sindels.

Particularly, by leveraging DanQ, which is a deep learning approach that can predict thousands of epigenomic profiles simultaneously given genomic sequence, we obtain 919 predicted tissue/cell type-specific epigenomic profiles for each nc-sindel. Since the alternative allele is assumed to disrupt the normal sequence pattern, which may cause the change of epigenomic profile and indicate the functional consequence of the nc-sindel, we also modify the reference sequence into alternative sequence by incorporating the nc-sindel by shifting the sequence upstream for insertion and downstream for deletion to make the length of the alternative sequence a fixed length of 1000bp. Accordingly, we also obtain another 919 epigenomic profiles using the alternative sequence.

To demonstrate the necessity to include both CADD functional annotations and multi-omics features derived from DanQ, we use five-fold cross-validation on 44 tissues in GTEx to evaluate three feature sets of TIVAN-indel: (i) 45 CADD functional annotations; (ii) 45 CADD functional annotations+919 multi-omics features under reference sequence; (iii) 45 CADD functional annotations+919 multi-omics features under both reference and alternative sequence. We report the AUROC and AUPRC for each tissue and use the median of AUROC and AUPRC across 44 tissues to compare the prediction performance among three feature sets.

As a result (Figure 2, S1), we find that inclusion of multi-omics features under the reference sequence will slightly improve both the median AUROC (CADD annotations:0.858, CADD annotations+multi-omics features (ref seq):0.859) and the median AUPRC (CADD annotations:0.843, CADD annotations+multi-omics features (ref seq): 0.845). This observation indicates that adding more tissue/cell-type specific multi-omics features will improve the prediction performance as a nc-sindel regulates gene expression in a tissue/cell type-specific manner. Importantly, further adding multi-omics features under the alternative sequence (CADD annotations+multi-omics features (ref seq+alt seq)), the prediction performance is significantly improved by achieving the highest median AUROC (0.982) and highest median AUPRC (0.984). Indeed, it is the comparison between the reference sequence and alternative sequence, as well as the associated multi-omics features, that characterizes the functional consequence of a nc-sindel. Biologically, a nc-sindel can disrupt the normal sequence pattern, which can abnormally up-regulate or down-regulate the nearby gene expression. The disruption can be due to the loss/change of a present motif sequence or gain of a new motif sequence, which will reject or recruit the binding of transcription factors that are essential in the gene regulation. In the following sections, we will use all three feature sets as the default setting for TIVAN-indel.

**Figure 2:**
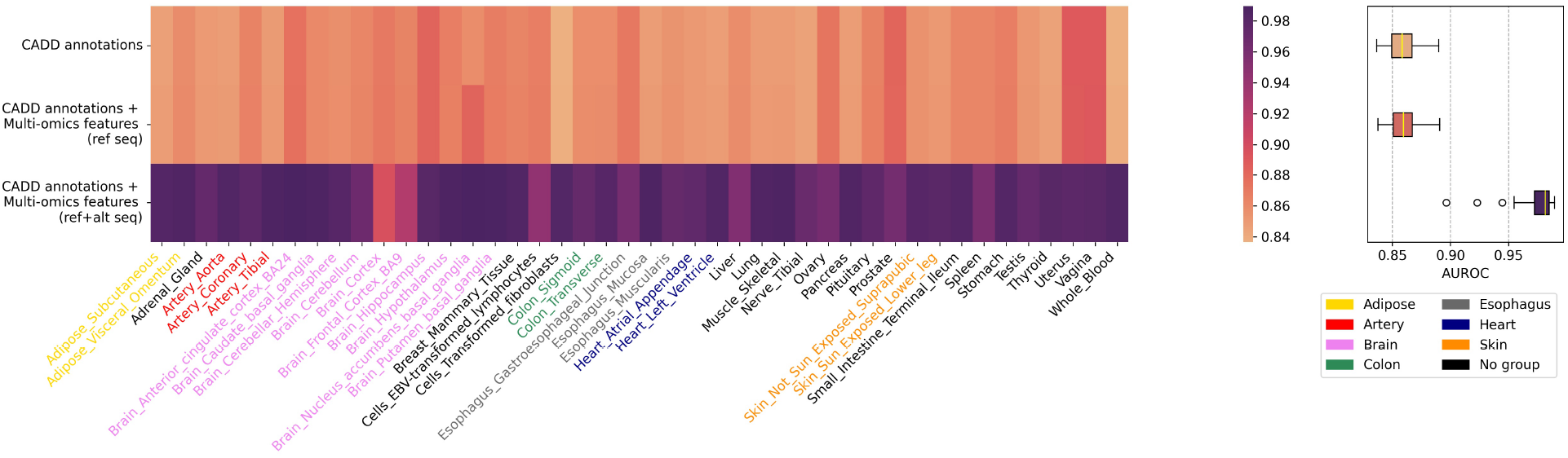
Comparison of three feature sets of TIVAN-indel: (i) 45 CADD functional annotations; (ii) 45 CADD functional annotations+919 multi-omics features under the reference sequence; (iii) 45 CADD functional annotations+919 multi-omics features under both the reference and alternative sequence across 44 tissues in GTEx. The median of AUROC across 44 tissues is reported for each feature set.

### 3.2 Evaluating TIVAN-indel to predict tissue-specific regulatory noncoding small indels using a cross-validation within-tissue approach

We adopt the cross-validation within-tissue approach to evaluate TIVAN-indel and the competing methods using labeled nc-sindels from 44 tissues in GTEx. For TIVAN-indel, we adopt five-fold cross-validation to obtain the prediction score for each fold of the five-folds for each GTEx tissue. The prediction scores of the five-folds are concatenated and compared to the true labels. For CADD and FATHMM-indel, we obtain the genome-wide prediction scores for sindels from CADD web server (https://cadd.gs.washington.edu/score) and FATHMM-indel web server (http://indels.biocompute.org.uk/). The precomputed scores are compared to the true labels directly. As a result, we find TIVAN-indel performs much better than CADD and FATHMM-indel for each tissue (Figure S2, S3).

However, the above comparison may be biased. Though the testing sets are labeled nc-sindels from GTEx, the training sets of the three methods are different, where TIVAN-indel uses labeled nc-sindels from the same tissue in GTEx, CADD adopts high-frequency human-derived alleles and FATHMM-indel utilizes pathogenic nc-sindels from HGMD. The different the training set makes CADD and FATHMM-indel focus on predicting the pathogenicity or deleteriousness of nc-sindels rather than their molecular impact in gene expression. To have a fair comparison and make CADD and FATHMM-indel focus on predicting the regulatory potential of nc-sindels, we retrain CADD and FATHMM-indel using five-fold cross-validation with four-folds of labeled nc-sindels as the training set and one-fold as the testing set for each GTEx tissue. Since both CADD and FATHMM-indel adopt SVM and share a similar feature set, we use Support Vector Machine (SVM) to train a classifier and use CADD functional annotations as the training features. We denote the trained SVM as “CADD/FATHMM-indel (SVM)”, which serves as the representative for both CADD and FATHMM-indel. In addition, we implement the common approach for deep learning models to predict functional noncoding variants. Specifically, we take predicted chromatin features for both reference allele and alternative alleles, and calculate the absolute difference features (919 features) and relative different features (919 features) [12], which are treated as training features in a boosted logistic regression model. We denote this approach as “DanQ/DeepSEA (Boosted LR)” and use five-fold cross-validation for performance evaluation.

Consequently, we find that TIVAN-indel outperforms CADD/FATHMM-indel (SVM) and DanQ/DeepSEA (Boosted LR) (Figure 3, S4) by achieving the highest AUROC and AUPRC (AUROC: 0.983 vs 0.784 vs 0.753; AUPRC: 0.986 vs 0.840 vs 0.816). The significant improvement of TIVAN-indel may be attributed to two factors. First, XGBoost, adopted by TIVAN-indel, has been demonstrated to be more powerful than conventional logistic regression [25] and SVM [26]. Second, TIVAN-indel considers both CADD functional annotations and tissue/cell type-specific multi-omics features. In contrast, CADD/FATHMM-indel (SVM) only considers CADD functional annotations, which mainly contains generic features such as conservation score, GC content and chromatin states. DanQ/DeepSEA (Boosted LR) mainly considers tissue/cell typespecific multi-omics features. Therefore, TIVAN-indel leverages the strength of both generic features and tissue/cell type-specific multi-omics features, which leads to the improvement of prediction performance.

**Figure 3:**
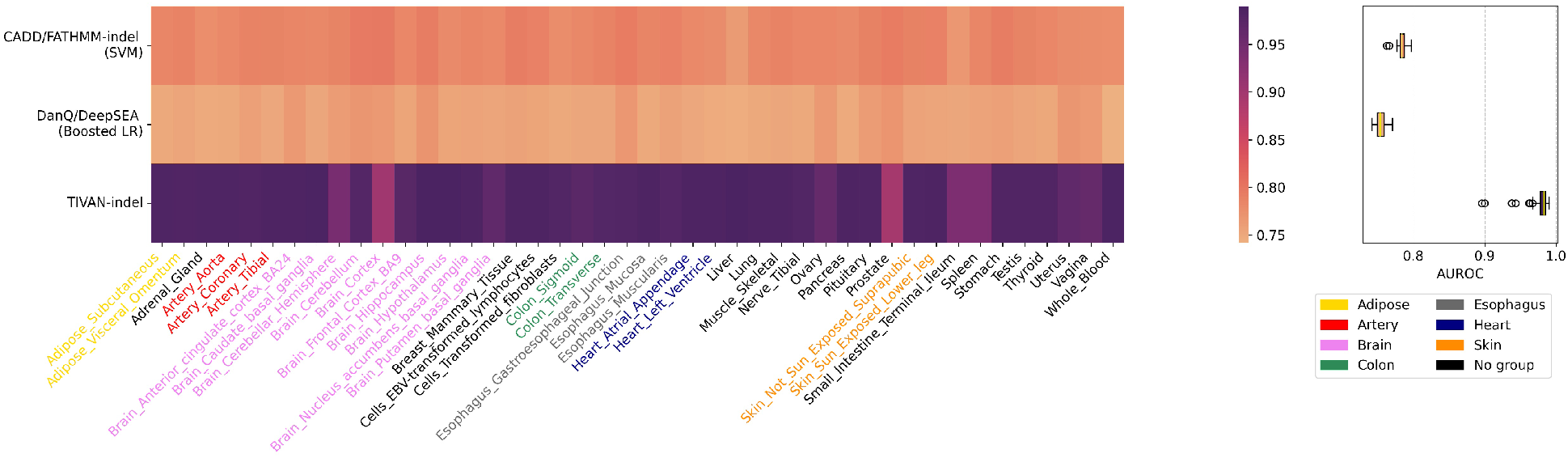
Comparison between TIVAN-indel, CADD/FATHMM-indel (SVM) and DanQ/DeepSEA (Boosted LR) using the within-tissue approach for 44 tissues/cell types in GTEx, where each method is trained and tested using five-fold cross-validation. AUROC is reported for each tissue/cell type.

### 3.3 Evaluating TIVAN-indel for predicting noncoding regulatory small indels using an independent cross-tissue approach

We further perform an independent cross-tissue approach, where all methods are trained using samples from one tissue and tested using samples from another tissue, to benchmark TIVAN-indel against the competing approaches for two reasons, First, the cross-tissue approach is an independent evaluation, where the training set and testing set are from different context. Therefore, this evaluation provides a more objective evaluation for all compared methods. Second, cross-tissue evaluation can identify which tissue is more powerful to predict regulatory nc-sindels from other tissues. Practically, we can use the TIVAN-indel model trained by the best-performed tissue identified in the cross-tissue prediction when there is a lack of training samples for the tissue in the testing set.

Accordingly, we train TIVAN-indel, CADD/FATHMM-indel (SVM) and DanQ/DeepSEA (Boosted LR) using one tissue/cell type and test the model on the remaining 43 tissues/cell types. The overlapped samples between the training set and the testing set are removed from the testing set. Consequently, we order the median of AUROC and AUPRC in 44 tissues/cell types (Figure 4, S5) and have three important findings. First, TIVAN-indel outperforms CADD/FATHMM-indel (SVM) and DanQ/DeepSEA (Boosted LR) (AUROC: 0.971 vs 0.831 vs 0.813; AUPRC: 0.964 vs 0.774 vs 0.722), which demonstrates the superiority of TIVAN-indel in the independent test setting. Second, within-tissue prediction achieves overall better performance than cross-tissue prediction for all methods due to the context-matching between training and testing set. The advantage of within-tissue prediction compared to cross-tissue prediction is more evident for CADD/FATHMM-indel (SVM) and DanQ/DeepSEA (Boosted LR). Last but most interestingly, we find the top-performed tissues are from the Brain category, which indicates that it is confident to use the TIVAN-indel model trained from the Brain category to predict regulatory nc-sindels from other tissues where the training samples for the targeted tissues are few or not readily available.

**Figure 4:**
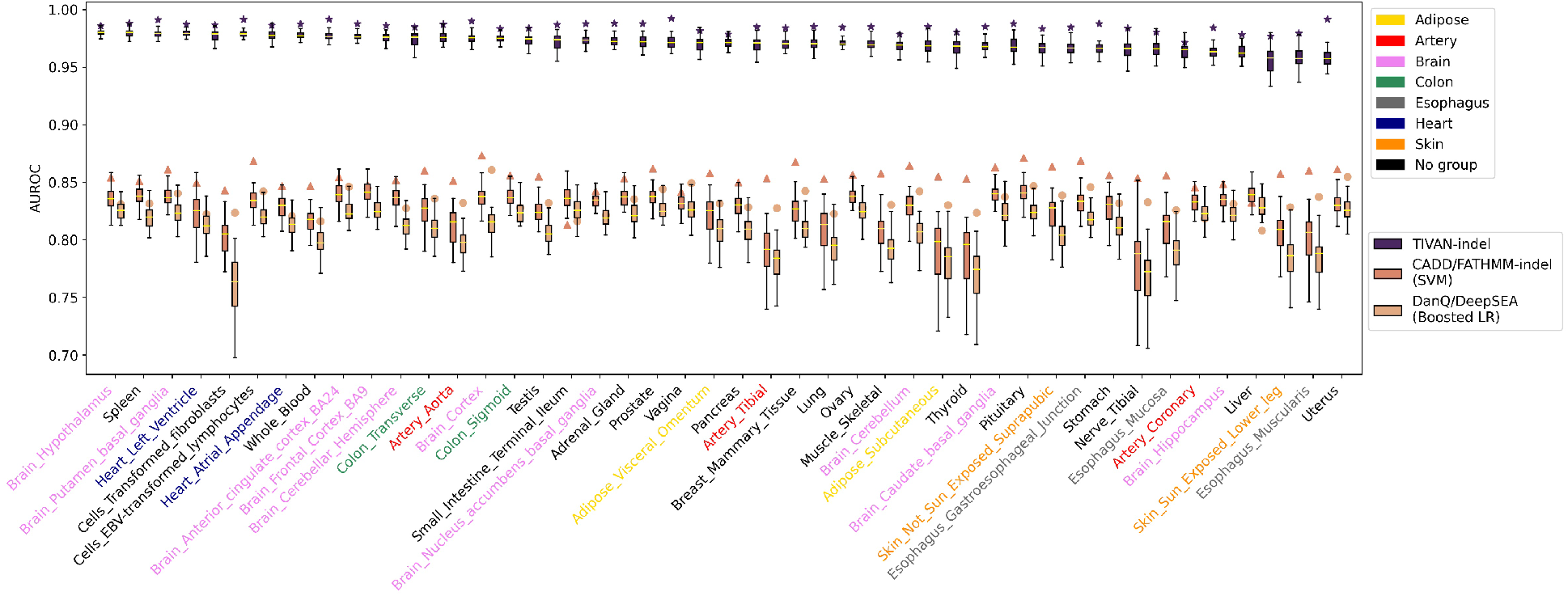
Comparison between TIVAN-indel, CADD/FATHMM-indel (SVM) and DanQ/DeepSEA (Boosted LR) using the independent cross-tissue approach for 44 tissues/cell types in GTEx, where each method is trained using one tissue/cell type and tested on the remaining 43 tissues/cell types. The overlapped samples between the training set and the testing set are removed from the testing set. For each tissue/cell type, AUROC is reported for the testing 43 tissue/cell types as demonstrated in the boxplot. The asterisk denotes the AUROC calculated from the cross-validation within-tissue approach. 44 tissues/cell types are colored in 8 tissue/cell types class.

### 3.4 Evaluating TIVAN-indel for predicting noncoding regulatory small indels in an independent study DICE

We have demonstrated that TIVAN-indel outperforms competing approaches based on the evaluation on GTEx data using both within-tissue and cross-tissue approaches. We further perform an independent evaluation, where the model is trained on GTEx data and tested on another QTL study named DICE (Database of Immune Cell Expression, Expression quantitative trait loci (eQTLs) and Epigenomics) (https://dice-database.org/) [27]. To create the testing set, we collect cell-type specific regulatory nc-sindels from 15 immune cell types in DICE, which include Classic and Non-classic monocytes, NK cells, Native B Cells, Naive/Stimulated CD4+ T cells, Naive/Stimulated CD8+ T cells, Naive/Memory Treg cells, Th1 cells, Th17 cells, Th1/17 cells, Th2 cells and Tfh. We use the same data processing steps for GTEx data to obtain the nc-sindels and create the positive and negative sets for each immune cell type. The number of regulatory nc-sindels for the 15 immune cel types are summarized in Table S5. Because immune cell types can be found in peripheral blood, we use the model trained from tissue “Whole blood” in GTEx data to predict the regulatory nc-sindels of 15 immune cell types in DICE.

Similarly, we find that TIVAN-indel consistently performs better than CADD/FATHMM-indel (SVM) and DanQ/DeepSEA (Boosted LR) by achieving highest AUC (0.973 vs 0.773 vs 0.744) and AUPRC (0.978 vs 0.792 vs 0.733) (Figure 5), S6). The high prediction accuracy from all methods indicates the biological connection between whole blood and immune cell types. Moreover, the independent evaluation furthers strengthens the superiority of using TIVAN-indel to predict the regulatory potential of nc-sindels.

**Figure 5:**
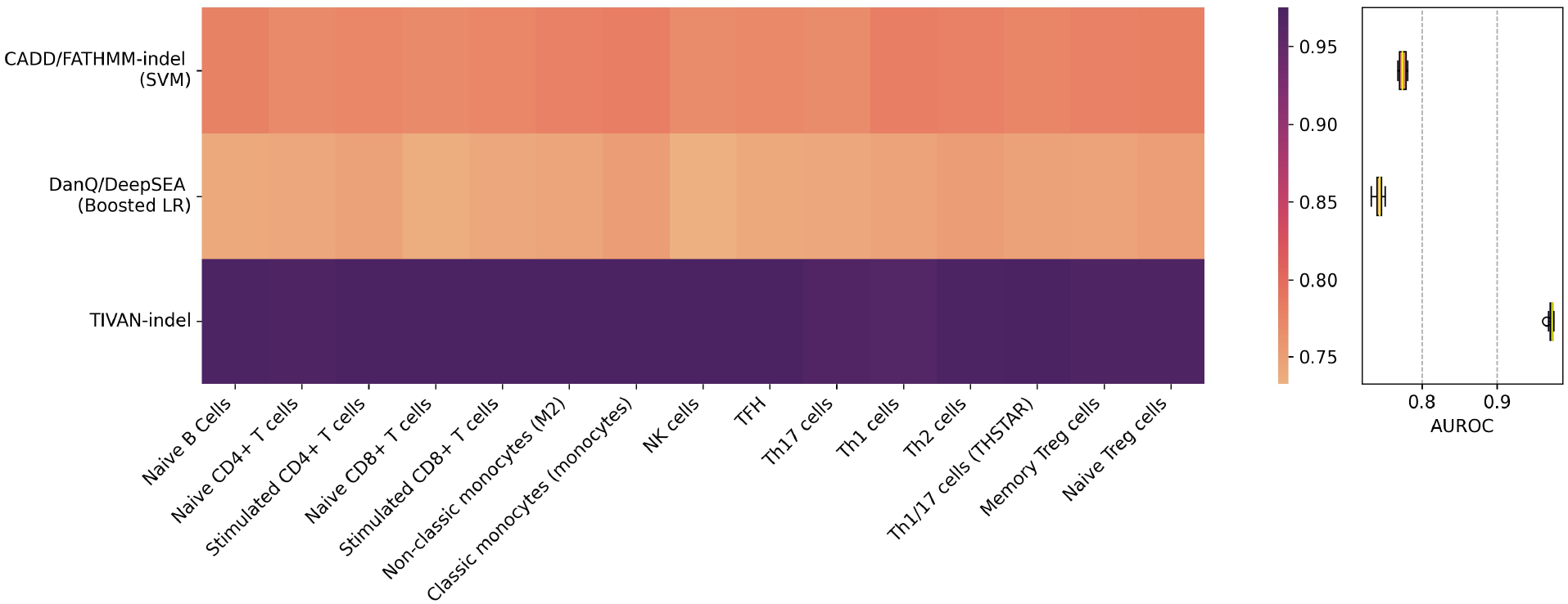
Comparison between TIVAN-indel, CADD/FATHMM-indel (SVM) and DanQ/DeepSEA (Boosted LR) by training on tissue “Whole blood” in GTEx data and testing on the regulatory nc-sindels of 15 immune cell types in DICE. The overlapped samples between training and testing sets are removed from the testing set. The AUROC is reported for each cell type.

### 3.5 Evaluating TIVAN-indel for predicting pathogenic noncoding small indels

Though the purpose for TIVAN-indel is to predict nc-indels that affect gene expression, we hereby evaluate the performance of TIVAN-indel in classifying pathogenic nc-sindels from the benign ones, which is the main task for CADD and FATHMM-indel. To do this, we download the latest version of “variant_summary.txt.gz” file from clinvar ftp website (https://ftp.ncbi.nlm.nih.gov/pub/clinvar/tab_delimited/). We collect small indels in the noncoding regions (hg19/GRCh37), which are annotated with “Insertion”, “Deletion”, “Indel” or “Duplication” and have both reference allele and alternative allele smaller than 100bp. We define pathogenic and benign nc-sindels with “ClinicalSignificance” noted as “Pathogenic” and “Benign”. We further select highly confident nc-sindels with the “Review status” noted as “multiple submitters, no conflicts” or “reviewed by expert panel”. Finally, we have 132 positive nc-sindels and 1104 negative nc-sindels

Similar to other work [11], we also adopt five-fold cross validation to evaluate all methods. We find that TIVAN-indel still outperforms CADD/FATHMM-indel (SVM) and DanQ/DeepSEA (Boosted LR) by achieving highest AUROC (Figure 6) and AUPRC (Figure S7). The observation indicates the robustness of TIVAN-indel, which can accurately predict both regulatory nc-sindels that regulate gene expression and nc-sindels that have a clinic impact.

**Figure 6:**
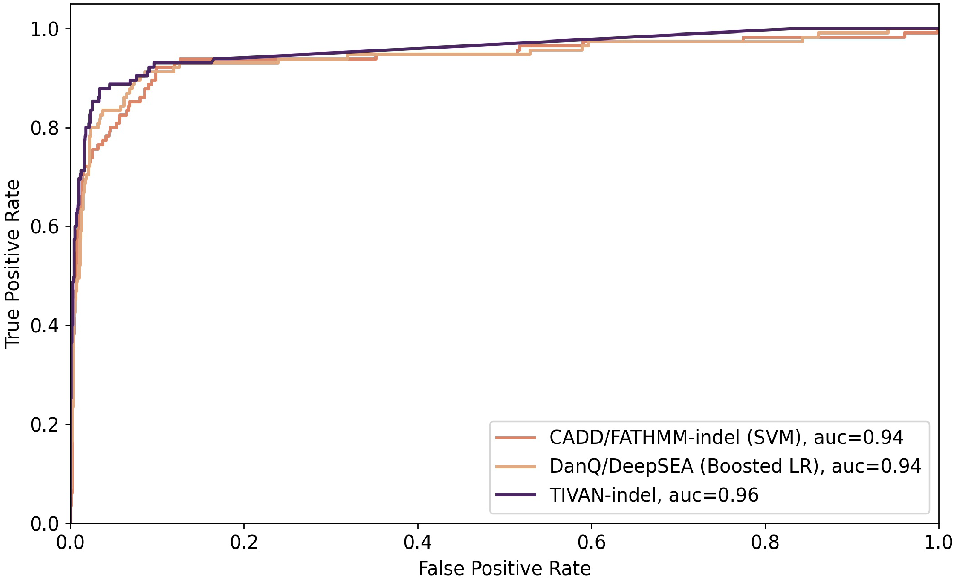
Comparison between TIVAN-indel, CADD/FATHMM-indel (SVM) and DanQ/DeepSEA (Boosted LR) on classifying pathogenic and benign nc-sindels in ClinVar. The AUROC is reported.

### 3.6 Enrichment analysis of regulatory nc-sindels in chromatin interactions and epigenomic regions

We hypothesize that regulatory nc-sindels need to be enriched in key regulatory regions such as enhancer, promoter and chromatin interactions in order to have an impact on gene expression. Moreover, we assume a good prediction of regulatory nc-sindels should achieve a similar or better enrichment in regulatory regions as the true labeled nc-sindels. Therefore, we perform the enrichment analysis for both true and predicted nc-sindels.

To obtain key regulatory regions, we collect tissue-specific promoter-centered capture Hi-C data across across 27 human tissue/cell types from 3DIV [20] and ChIP-seq data profiling open chromatin and key histone marks from Roadmap Epigenomics [22]. For Hi-C data, we create tissue-specific positive/negative promoter-enhancer (PE) and promoter-promoter (PP) chromatin interactions as our previous work [21]. For ChIP-seq data, we treat peak regions with the FDR less than 0.05 as the positive set and randomly sampled regions with a matched GC content distribution as the negative set. More details of the data collection and processing can be found in the Method section. We perform the enrichment analysis in a tissue-specific way by matching the tissue/cell type in GTEx and tissue/cell type of Hi-C data and ChIP-seq data.

In the enrichment analysis, we first calculate the number of positive/negative nc-sindels overlapped in positive/negative regulatory regions, which include chromatin interactions or peak regions respectively. For peak regions, we directly overlap genomic location of nc-sindels and the peak regions. For chromatin interactions, we overlap the genomic location of nc-sindels and the two anchors of each interaction. As long as the nc-sindel hits one of the two anchors, we consider the nc-sindel is overlapped with the interaction. Based on this calculation, we obtain *n*_11_: number of positive nc-sindels overlapped in positive regulatory regions; *n*_12_: number of positive nc-sindels overlapped in negative regulatory regions; *n*_21_: number of negative nc-sindels overlapped in positive regulatory regions; *n*_22_: number of negative nc-sindels overlapped in negative regulatory regions. Then, we perform a fisher’s exact test to calculate the enrichment and report pvalues by assuming positive nc-sindels is more enriched in positive regulatory regions. The enrichment is defined as the odds ratio in the formula as 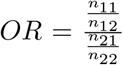. Moreover, we use a threshold of 0.5 to classify the prediction into predicted positive/negative nc-sindels, and calculate the enrichment using predicted labels and compared to the enrichment calculated using true labels.

For chromatin interaction, we perform the enrichment analysis of both true and predicted regulatory nc-sindels from GTEx in tissue-matching PE and PP from 3DIV respectively (Figure 7A, B). We find that both true and predicted labels are enriched in PE across 20 tissues/cell types among matched 24 tissues/cell types (OR>1). Similarly, both true and predicted labels are enriched in PP across 16 tissues/cell types among 19 matched tissues/cell types (OR>1). This observation indicates that regulatory nc-sindels are more likely to be enriched in PE and PP in the same tissue/cell type. Moreover, the enrichment patterns for the true labels and predicted labels are similar and the overall enrichment of predicted labels is slightly lower than the true labels. For PE, both true and predicted labels are significantly enriched in top-ranked tissues/cell types such as “Small Intestine Terminal Ileum” (OR=2.77 and pvalue=2.74 × 10^-5^ for true labels; OR=2.13 and pvalue=0.002 for predicted labels), “Stomach” (OR=1.86 and pvalue=1.15 × 10^-9^ for true labels, OR=1.58 and pvalue=7.97 × 10^-6^ for predicted labels) and “Lung” (OR=1.47 and pvalue=8.79 × 10^-7^ for true labels; OR=1.48; pvalue=9.39 × 10^-7^ for predicted labels). In contrast, true labels but not predicted labels are significantly enriched in “Ovary” (OR=1.59 and pvalue=0.035 for true labels; OR=1.34 and pvalue=0.19 for predicted labels). For PP, true labels are more enriched in “Ovary” (OR=5.58 and pvalue=0.002), “Pancreas” (OR=2.48 and pvalue=2.10 × 10^-5^), “Liver” (OR=2.18 and pvalue=0.005) and Artery Coronary (OR=2.12; pvalue=6.92 × 10^-4^). Predicted labels also keep a similar enrichment trend though the enrichment in “Ovary is much lower (OR=5.58 for true labels vs OR=1.83 for predicted labels). Interestingly, we find true labels and predicted labels are enriched in “Ovary” for both PE and PP.

**Figure 7:**
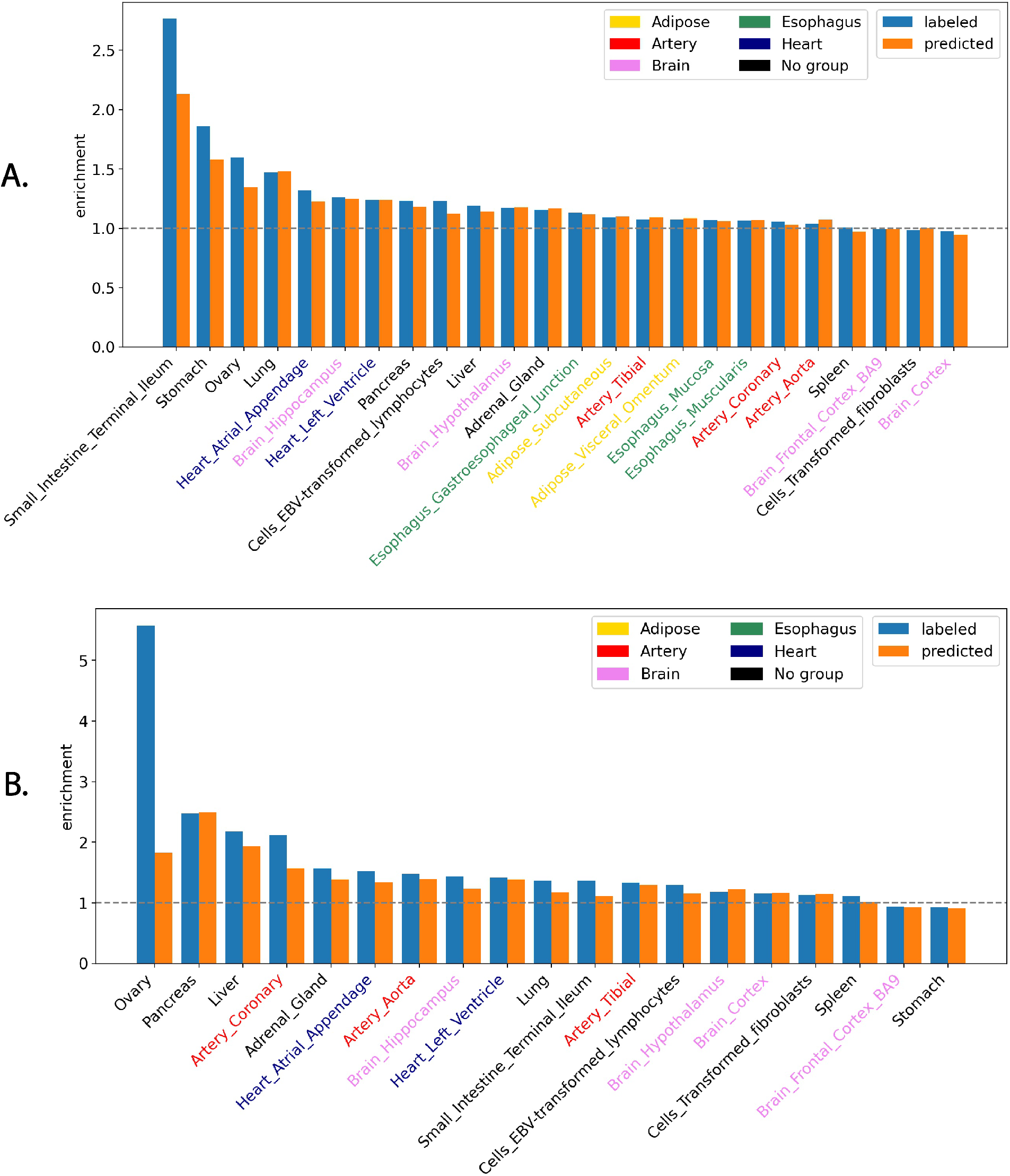
Enrichment analysis of both labeled and predicted regulatory nc-sindels from GTEx in tissuematching pcHi-C data from 3DIV.

Similarly, we perform the enrichment analysis for both labeled and predicted regulatory nc-sindels from GTEx in tissue-matching epigenomic regions from Roadmap Epigenomics, which include open chromatin, five active histone marks and two repressive histone marks (Figure 8A-H). Consequently, we find that both and predicted labels are enriched in open chromatins (OR>1) and achieve comparable enrichment across all tissues/cell types (Figure 8A). Similarly, both true and predicted labels are significantly enriched in H3K36me3 associated with actively transcribed genes across all tissues/cell types (OR>1 and pvalue<0.05) (Figure 8B). Importantly, we find that predicted labels show a higher enrichment than true labels for all tissues/cell types (median OR=3.71 for predicted labels vs median OR=2.62 for true labels). We define the true labels based on statistical significance but the significant nc-sindels are not necessarily the causal ones. Therefore, the higher enrichment of predicted labels than true labels indicates that TIVAN-indel can accurately predict the regulatory nc-sindels.

**Figure 8:**
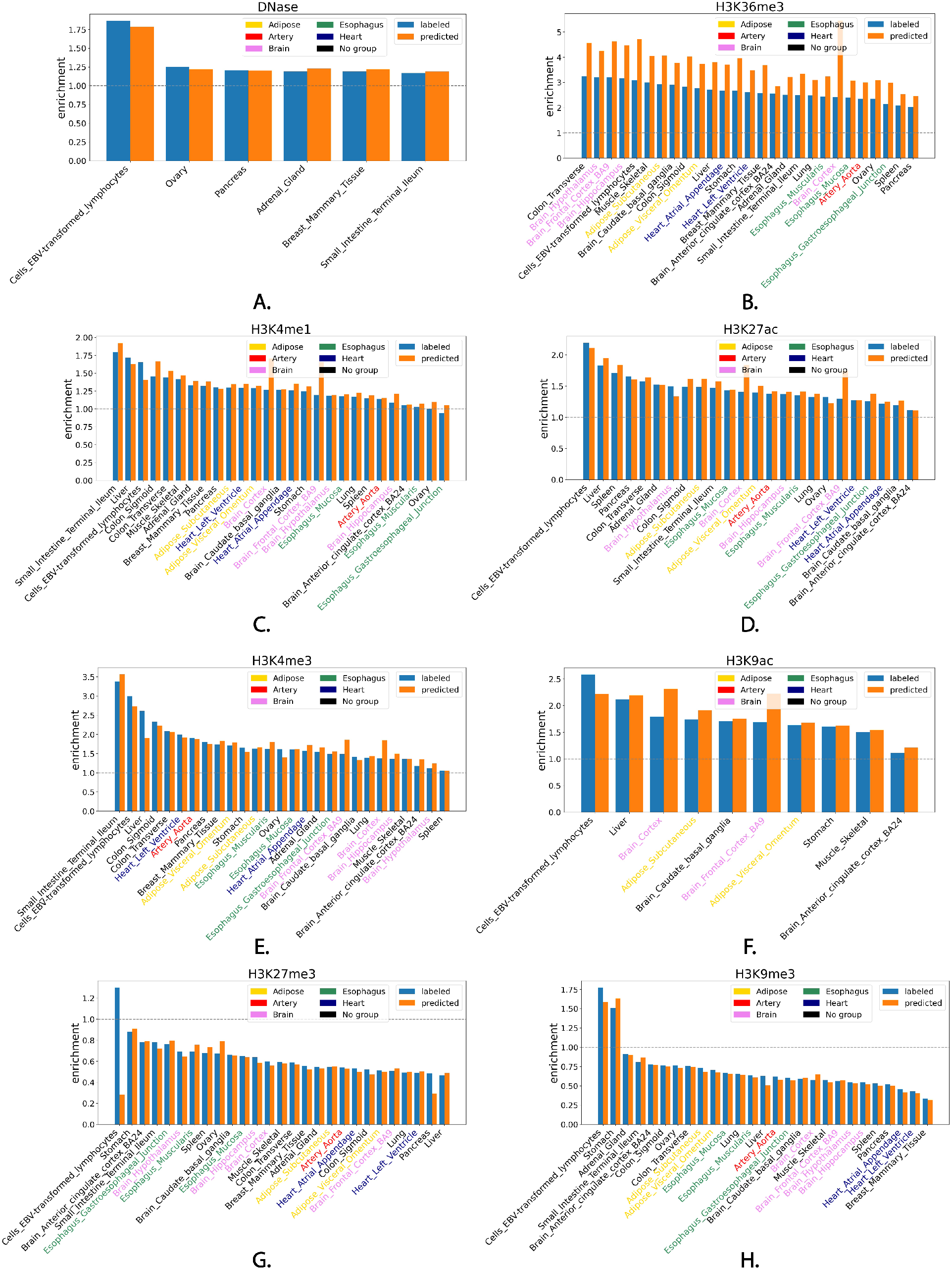
Enrichment analysis of labeled and predicted regulatory nc-sindels from GTEx in tissue-matching ChIP-seq data from Roadmap Epigenomics.

Moreover, both true labels and predicted labels are enriched in H3K4me1 and H3K27ac, which are histone marks associated with enhancer activation, across most tissues/cell types in a similar enrichment trend (OR>1) (Figure 8C,D). Interestingly, we find that the enrichment of predicted labels is higher than true labels in “Brain Cortex” and “Brain Frontal Cortex BA9” for both H3K4me1 (OR=1.71 and pvalue=4.75 × 10^-11^ for predicted labels vs OR=1.27 and pvalue=3.45 × 10^-3^ for true labels in “Brain Cortex”; OR=1.66 and pvalue=1.22 × 10^-7^ for predicted labels vs OR=1.20 and pvalue=6.02 × 10^-2^ for true labels in “Brain Frontal Cortex BA9”) and H3K27ac (OR=1.84 and pvalue=6.35 × 10^-13^ for predicted labels vs OR=1.40 and pvalue=5.96 × 10^-5^ for true labels in “Brain Cortex”; OR=1.75 and pvalue=3.24 × 10-8 for predicted labels vs OR=1.29 and pvalue=1.11 ×^1^ 0^-2^ for true labels in “Brain Frontal Cortex BA9”). Similar to active histone marks associated with enhancer activation, both true and predicted labels are enriched in H3K4me3 and H3K9ac, which are histone markers associated with promoter activation, across all matched tissues/cell types in a similar enrichment trend (OR>1) (Figure 8E,F). In addition, the enrichment of predicted labels is higher than true labels in “Brain Cortex” and “Brain Frontal Cortex BA9” for both H3K4me3 (OR=1.84 and pvalue=4.36 × 10^-6^ for predicted labels vs OR=1.38 and pvalue=1.73 × 10^-2^ for true labels in “Brain Cortex”; OR=1.86 and pvalue=5.11 × 10^-5^ for predicted labels vs OR=1.49 and pvalue=9.42 × 10^-3^ for true labels in “Brain Frontal Cortex BA9”) and H3K9ac (OR=2.32 and pvalue=2.67 × 10^-13^ for predicted labels vs OR=1.79 and pvalue=2.44 × 10^-7^ for true labels in “Brain Cortex”; OR=2.22 and pvalue=8.39 × 10^-10^ for predicted labels and OR=1.69 and pvalue=5.08 × 10^-5^ in “Brain Frontal Cortex BA9”). Again, the higher enrichment of predicted labels than true labels in histone marks associated in both promoter activation and enhancer activation indicates that TIVAN-indel can accurately predict the regulatory nc-sindels in “Brain Cortex” and “Brain Frontal Cortex BA9”.

In contrast, we find that both true labels and predicted labels are depleted in repressive chromatin marks such as H3K27me3 and H3K9me3 in most tissue/cell types (OR<1) (Figure 8G,H). For H3K9me3, both predicted and true labels are depleted in all tissues/cell types except for “Cells EBV-transformed lymphocytes” and “Stomach”. For H3K27me3, predicted labels are depleted while true labels are enriched in “Cells EBV-transformed lymphocytes” (OR=0.28 for predicted labels vs OR=1.30 for true labels). The depletion instead of enrichment for predicted labels indicates that TIVAN-indel can improve the prediction for regulatory nc-sindels in “Cells EBV-transformed lymphocytes”.

## Conclusion

We develop TIVAN-indel, which is a genome-wide variant annotation tool that assigns higher scores to nc-sindels that are more likely to alter the expression levels of nearby genes. In contrast to existing approaches such as CADD and FATHMM-indel, which focus on pathogenicity or deleteriousness, TIVAN-indel can interpret nc-indels’ molecular impact in gene expression and thus can offer a more comprehensive mechanistic understanding of the functional role of nc-indels in disease pathogenesis. Importantly, TIVAN-indel is a complementary tool to identify regulatory nc-indels considering the limitation of current sequencing experiments such as the small sample size, low minor allele frequencies (MAFs), small effects or lack of matched gene expression data for the genotype data.

The advantages of TIVAN-indel lies on two main aspects. First, it leverages a comprehensive set of tissue/cell type-specific nc-sindels from GTEx data to train a tissue/cell type-specific model. This is important because gene expression varies in different tissues and a nc-sindel may affect the nearby gene expression in a tissue/cell type-specific manner. Second, TIVAN-indel utilizes a rich feature set, which includes both CADD functional annotations and multi-omics features that are derived from DanQ, a sequence-based deep learning model. The expanded feature set will allow both generic and context-specific characterization of possible mechanistic insights of how a nc-sindel impacts the gene expression under reference allele and alternative allele, which in turn improve the prediction for regulatory nc-sindels. By using GTEx data, we demonstrates that using both feature sets improve the prediction performance compared to using either feature set.

We adopt both within-tissue and cross-tissue approaches to evaluate TIVAN-indel and its competing methods. In the within-tissue approach, we use five-fold cross validation to evaluate the prediction within the same tissue. In the cross-tissue approach, we train the model using one tissue and evaluate the model in the remaining tissues. Since existing approaches focus on evaluating the pathogenicity or deleteriousness of nc-sindels, we retrain CADD, FATHMM-indel and DeepSEA/DanQ using the same training set as TIVAN-indel, which are labeled nc-sindels from GTEx, to allow a fair comparison. Consequently, we find that TIVAN-indel still significantly outperforms existing computational tools in both ways of evaluation. To further demonstrate the robustness of TIVAN-indel, we train TIVAN-indel along with its competing methods using “Whole Blood” tissue from GTEx data and evaluate all methods using 15 immune cell types from an independent study DICE. This independent evaluation further confirms the superiority of TIVAN-indel in predicting regulatory nc-indels.

Moreover, we perform an enrichment analysis for nc-sindels by hypothesizing that regulatory nc-sindels need to be enriched in key regulatory regions such as enhancer, promoter and chromatin interactions in order to have an impact on gene expression. As expected, we find that both true and predicted labels are enriched in promoter-promoter and promoter-enhancer chromatin interactions across most tissues/cell types. Similarly, both true and predicted labels are enriched in histone marks associated with open chromatin, promoter, enhancer and gene activation. In contrast, they are depleted in repressive histone marks associated with downregulation of nearby genes. Importantly, both true and predicted labels share a similar the enrichment pattern in open chromatin, active histone marks and repressive histone marks, which indicates a good prediction made by TIVAN-indel. Interestingly, the enrichment of predicted labels is higher than true labels for active histone marks associated with promoter and enhancer activation in “Brain Cortex” and “Brain Frontal Cortex BA9”. Similarly, predicted labels are depleted while true labels are enriched in “Cells EBV-transformed lymphocytes” for H3K27me3. These observations indicate that TIVAN-indel can accurately predict regulatory nc-sindels, which are otherwise missed by the measurement of statistical significance.

We deliver TIVAN-indel as an open-source Python toolkit, which allows to train the model and predict the functional scores for given sc-indels. The functional scores can be used to prioritize regulatory nc-sindels, which can help narrow down the candidates for experimental validation by using Massively Parallel Reporter Assay (MPRA) or CRISPR-Cas9. This application makes TIVAN-indel practically useful considering current high-throughput sequencing technologies are still infeasible to validate all nc-sindels in human genome due to technical and financial challenges.

In this work, TIVAN-indel focus on predicting nc-sindels that regulate gene expression by leveraging GTEx data, which offers matched gene expression data and genotype data across multiple tissues and cell types. However, given the readily availability of matched genotype data and other types of functional omics data, TIVAN-indel can be easily extended to predict nc-sindels that impact on molecular phenotypes such as chromatin accessibility, DNA methylation and transcription factor binding. Therefore, multiple sets of functional scores for measuring the functional potential of nc-sindels in different aspects of molecular functions can be integrated to better annotate and characterize the functional role of nc-sindels, which can provide a comprehensive functional landscape for nc-sindels in human genome.

## Acknowledgement

This work was supported by National Institute of General Medical Sciences of the National Institutes of Health under Award Number R35GM142701 to LC.

