## Supplementary material for "TIVAN-indel: A computational framework for annotating and predicting noncoding regulatory small insertion and deletion"

#### 1 Supplementary tables

Table S1: Summary of the number of labeled nc-sindels in 44 tissues/cell types in GTEx

| Tissue | #Samples |
|---|---|
| Adipose Subcutaneous | 39524 |
| Adipose Visceral Omentum | 16724 |
| Adrenal Gland | 11052 |
| Artery Aorta | 26777 |
| Artery Coronary | 7649 |
| Artery Tibial | 39095 |
| Brain Anterior cingulate cortex BA24 | 3226 |
| Brain Caudate basal ganglia | 7231 |
| Brain Cerebellar Hemisphere | 10148 |
| Brain Cerebellum | 13809 |
| Brain Cortex | 7540 |
| Brain Frontal Cortex BA9 | 5728 |
| Brain Hippocampus | 3021 |
| Brain Hypothalamus | 3135 |
| Brain Nucleus accumbens basal ganglia | 5863 |
| Brain Putamen basal ganglia | 4358 |
| Breast Mammary Tissue | 15731 |
| Cells EBV-transformed lymphocytes | 10236 |
| Cells Transformed fibroblasts | 43632 |
| Colon Sigmoid | 9972 |
| Colon Transverse | 16671 |
| Esophagus Gastroesophageal Junction | 10225 |
| Esophagus Mucosa | 33910 |
| Esophagus Muscularis | 31487 |
| Heart Atrial Appendage | 15059 |
| Heart Left Ventricle | 18523 |
| Liver | 4978 |
| Lung | 31046 |
| Muscle Skeletal | 36293 |
| Nerve Tibial | 45474 |
| Ovary | 4349 |
| Pancreas | 16630 |
| Pituitary | 6337 |
| Prostate | 4172 |
| Skin Not Sun Exposed Suprapubic | 21141 |
| Skin Sun Exposed Lower leg | 39416 |
| Small Intestine Terminal Ileum | 3289 |
| Spleen | 7860 |
| Stomach | 13390 |
| Testis | 30251 |
| Thyroid | 45561 |
| Uterus | 2258 |
| Vagina | 2390 |
| Whole Blood | 33580 |

#### Table S2: Summary of CADD annotations used for TIVAN-indel

| | Annotation | Annotation Class | Annotation Type |
|---|---|---|---|
| 1 | Consequence | VEP consequence, priority selected by potential impact | categorical |
| 2 | ConsScore | Custom deleterious score assigned to Consequence | numeric |
| 3 | GC | Percent GC in a window of $+/-$ 75bp | numeric |
| 4 | CpG | Percent CpG in a window of $+/-$ 75bp | numeric |
| 5 | minDistTSS | Distance to closest Transcribed Sequence Start (TSS) | numeric |
| 7 | minDistTSE | Distance to closest Transcribed Sequence Start (TSE) | numeric |
| 7 | priPhCons | Primate PhastCons conservation score | numeric |
| 8 | mamPhCons | Mammalian PhastCons conservation score | numeric |
| 9 | verPhCons | Vertebrate PhastCons conservation score | numeric |
| 10 | priPhyloP | Primate PhyloP score | numeric |
| 11 | mamPhyloP | Mammalian PhyloP score | numeric |
| 12 | verPhyloP | Vertebrate PhyloP score | numeric |
| 13 | bStatistic | Background selection score | numeric |
| 14 | cHmmTssA | Number of 48 cell types in chromHMM state | numeric |
| 15 | cHmmTssAFlnk | Number of 48 cell types in chromHMM state | numeric |
| 16 | cHmmTxFlnk | Number of 48 cell types in chromHMM state | numeric |
| 17 | cHmmTx | Number of 48 cell types in chromHMM state | numeric |
| 18 | cHmmTxWk | Number of 48 cell types in chromHMM state | numeric |
| 19 | cHmmEnhG | Number of 48 cell types in chromHMM state | numeric |
| 20 | cHmmEnh | Number of 48 cell types in chromHMM state | numeric |
| 21 | cHmmZnfRpts | Number of 48 cell types in chromHMM state | numeric |
| 22 | cHmmHet | Number of 48 cell types in chromHMM state | numeric |
| 23 | cHmmTssBiv | Number of 48 cell types in chromHMM state | numeric |
| 24 | cHmmBivFlnk | Number of 48 cell types in chromHMM state | numeric |
| 25 | cHmmEnhBiv | Number of 48 cell types in chromHMM state | numeric |
| 26 | cHmmReprPC | Number of 48 cell types in chromHMM state | numeric |
| 27 | cHmmReprPCWk | Number of 48 cell types in chromHMM state | numeric |
| 28 | cHmmQuies | Number of 48 cell types in chromHMM state | numeric |
| 29 | GerpN | Neutral evolution score defined by GERP++ | numeric |
| 30 | GerpS | Rejected Substitution score defined by GERP++ | numeric |
| 31 | EncH3K27Ac | Encode H3K27ac levels (from 14 cell lines) | numeric |
| 32 | EncH3K4Me1 | Encode H3K4me1 levels (from 14 cell lines) | numeric |
| 32 | EncH3K4Me3 | Encode H3K4me3 levels (from 14 cell lines) | numeric |
| 34 | EncNucleo | Encode nucleosome occupancy levels (from 14 cell lines) | numeric |
| 35 | Segway | transcriptome/epigenome (ENCODE/Roadmap) | categorical |
| 36 | Dist2Mutation | Distance between the closest BRAVO SNV up and downstream | numeric |
| 37 | Freq100bp | Number of frequent (MAF $>$0.05) BRAVO SNV in 100 bp window nearby | numeric |
| 38 | Rare100bp | Number of rare (MAF $<$ 0.05) BRAVO SNV in 100 bp window nearby | numeric |
| 39 | Sngl100bp | Number of single occurrence BRAVO SNV in 100 bp window nearby | numeric |
| 40 | Freq1000bp | Number of frequent (MAF $>$0.05) BRAVO SNV in 1000 bp window nearby | numeric |
| 41 | Rare1000bp | Number of rare (MAF $<$ 0.05) BRAVO SNV in 1000 bp window nearby | numeric |
| 42 | Sngl1000bp | Number of single occurrence BRAVO SNV in 1000 bp window nearby | numeric |
| 43 | Freq10000bp | Number of frequent (MAF $>$0.05) BRAVO SNV in 10000 bp window nearby | numeric |
| 44 | Rare10000bp | Number of rare (MAF $<$ 0.05) BRAVO SNV in 10000 bp window nearby | numeric |
| 45 | Sngl10000bp | Number of single occurrence BRAVO SNV in 10000 bp window nearby | numeric |

Table S3: Summary of pcHiC interactions

| Tissue | #Interactions | Tissue/cell type | #PE (FDR $< 0.1$) | #PP (FDR $< 0.1$) |
|---|---|---|---|---|
| Adipose Subcutaneous | 41363 | Fat | 56057 | $\times$ |
| Adipose Visceral Omentum | 17567 | Fat | 56057 | $\times$ |
| Adrenal Gland | 11626 | Adrenal Gland | 2556 | 5421 |
| Artery Aorta | 28031 | Aorta | 5224 | 788 |
| Artery Coronary | 8050 | Aorta | 5224 | 788 |
| Artery Tibial | 40984 | Aorta | 5224 | 788 |
| Brain Anterior cingulate cortex BA24 | 3425 | Dorsorateral prefrontal cortex | 26084 | 6224 |
| Brain Caudate basal ganglia | 7643 | Dorsorateral prefrontal cortex | 26084 | 6224 |
| Brain Cerebellar Hemisphere | 10685 | Dorsorateral prefrontal cortex | 26084 | 6224 |
| Brain Cerebellum | 14543 | Dorsorateral prefrontal cortex | 26084 | 6224 |
| Brain Cortex | 7973 | Dorsorateral prefrontal cortex | 26084 | 6224 |
| Brain Frontal Cortex BA9 | 6054 | Dorsorateral prefrontal cortex | 26084 | 6224 |
| Brain Hippocampus | 3201 | Hippocampus | 17434 | 2519 |
| Brain Hypothalamus | 3324 | Hippocampus | 17434 | 2519 |
| Brain Nucleus accumbens basal ganglia | 6183 | Dorsorateral prefrontal cortex | 26084 | 6224 |
| Brain Putamen basal ganglia | 4617 | Dorsorateral prefrontal cortex | 26084 | 6224 |
| Breast Mammary Tissue | 16546 | $\times$ | - | - |
| Cells EBV-transformed lymphocytes | 10756 | GM12878+GM19240 Lymphoblastoid Cell Line | 4903 | 4446 |
| Cells Transformed fibroblasts | 45580 | Fibroblast cells | 13217 | 4156 |
| Colon Sigmoid | 10518 | Sigmoid colon | $\times$ | 10338 |
| Colon Transverse | 17565 | Sigmoid colon | $\times$ | 10338 |
| Esophagus Gastroesophageal Junction | 10758 | Esophagus | 20049 | $\times$ |
| Esophagus Mucosa | 35559 | Esophagus | 20049 | $\times$ |
| Esophagus Muscularis | 33027 | Esophagus | 20049 | $\times$ |
| Heart Atrial Appendage | 15849 | Left Ventricle | 1565 | 968 |
| Heart Left Ventricle | 19472 | Left Ventricle | 1565 | 968 |
| Liver | 5252 | Liver | 5325 | 706 |
| Lung | 32528 | Lung | 1188 | 661 |
| Muscle Skeletal | 37990 | $\times$ | - | - |
| Nerve Tibial | 47555 | $\times$ | - | - |
| Ovary | 4586 | Ovary | 1105 | 187 |
| Pancreas | 17451 | Pancreas | 2594 | 396 |
| Pituitary | 6703 | $\times$ | - | - |
| Prostate | 4410 | $\times$ | - | - |
| Skin Not Sun Exposed Suprapubic | 22215 | $\times$ | - | - |
| Skin Sun Exposed Lower leg | 41294 | $\times$ | - | - |
| Small Intestine Terminal Ileum | 3500 | Small Bowel | 972 | 704 |
| Spleen | 8307 | Spleen | 22392 | 5866 |
| Stomach | 14100 | Gastric tissue | 1610 | 900 |
| Testis | 31732 | $\times$ | - | - |
| Thyroid | 47647 | $\times$ | - | - |
| Uterus | 2398 | $\times$ | - | - |
| Vagina | 2554 | $\times$ | - | - |
| Whole Blood | 35143 | $\times$ | - | - |

Table S4: Summary of ChIP-seq data in Roadmap Epigenomics

| Tissue | #Peaks | Tissue/cell type | DNase | H3K27me3 | H3K4me1 | H3K9ac | H3K27ac | H3K36me3 | H3K4me3 | H3K9me3 |
|---|---|---|---|---|---|---|---|---|---|---|
| Adipose Subcutaneous | 41363 | Adipose Nuclei | × | 187394 | 257141 | 98296 | 120547 | 252686 | 83253 | 208451 |
| Adipose Visceral Omentum | 17567 | Adipose Nuclei | × | 187394 | 257141 | 98296 | 120547 | 252686 | 83253 | 208451 |
| Adrenal Gland | 11626 | Fetal Adrenal Gland | 381299 | 287466 | 276138 | × | 155447 | 292480 | 33369 | 211821 |
| Artery Aorta | 28031 | Aorta | × | 47224 | 132931 | × | 130935 | 62301 | 37104 | 71996 |
| Artery Coronary | 8050 | × | - | - | - | - | - | - | - | - |
| Artery Tibial | 40984 | × | - | - | - | - | - | - | - | - |
| Brain Anterior cingulate cortex BA24 | 3425 | Brain Cingulate Gyrus | × | 71546 | 255504 | 125356 | 166221 | 198074 | 74743 | 89353 |
| Brain Caudate basal ganglia | 7643 | Brain Anterior Caudate | × | 37011 | 271201 | 166608 | 209550 | 224169 | 85705 | 149449 |
| Brain Cerebellar Hemisphere | 10685 | × | - | - | - | - | - | - | - | - |
| Brain Cerebellum | 14543 | × | - | - | - | - | - | - | - | - |
| Brain Cortex | 7973 | Brain Dorsolateral Prefrontal Cortex | × | 108957 | 218893 | 102626 | 198908 | 187780 | 71215 | 90299 |
| Brain Frontal Cortex BA9 | 6054 | Brain Dorsolateral Prefrontal Cortex | × | 108957 | 218893 | 102626 | 198908 | 187780 | 71215 | 90299 |
| Brain Hippocampus | 3201 | Brain Hippocampus Middle | × | 118969 | 224117 | × | 150278 | 189289 | 75240 | 124885 |
| Brain Hypothalamus | 3324 | Brain Hippocampus Middle | × | 118969 | 224117 | × | 150278 | 189289 | 75240 | 124885 |
| Brain Nucleus accumbens basal ganglia | 6183 | × | - | - | - | - | - | - | - | - |
| Brain Putamen basal ganglia | 4617 | × | - | - | - | - | - | - | - | - |
| Breast Mammary Tissue | 16546 | Breast vHMEC | 239398 | 121677 | 344219 | × | × | 212637 | 51855 | 188686 |
| Cells EBV-transformed lymphocytes | 10756 | Lymphoblastoid Cells | 224379 | 401 | 115050 | 47553 | 75909 | 127502 | 67368 | 27438 |
| Cells Transformed fibroblasts | 45580 | × | - | - | - | - | - | - | - | - |
| Colon Sigmoid | 10518 | Sigmoid Colon | × | 135997 | 110437 | × | 179713 | 279430 | 59431 | 111694 |
| Colon Transverse | 17565 | Sigmoid Colon | × | 135997 | 110437 | × | 179713 | 279430 | 59431 | 111694 |
| Esophagus Gastroesophageal Junction | 10758 | Esophagus | × | 50452 | 237103 | × | 149589 | 239923 | 50117 | 229209 |
| Esophagus Mucosa | 35559 | Esophagus | × | 50452 | 237103 | × | 149589 | 239923 | 50117 | 229209 |
| Esophagus Muscularis | 33027 | Esophagus | × | 50452 | 237103 | × | 149589 | 239923 | 50117 | 229209 |
| Heart Atrial Appendage | 15849 | Left Ventricle | × | 108042 | 254975 | × | 150090 | 271995 | 39466 | 156533 |
| Heart Left Ventricle | 19472 | Left Ventricle | × | 108042 | 254975 | × | 150090 | 271995 | 39466 | 156533 |
| Liver | 5252 | Liver | × | 86572 | 229385 | 102736 | 115620 | 296963 | 85967 | 249912 |
| Lung | 32528 | Lung | × | 49060 | 295061 | × | 213427 | 322694 | 76220 | 401423 |
| Muscle Skeletal | 37990 | Skeletal Muscle Male | × | 180943 | 234626 | 137343 | × | 181211 | 76955 | 65435 |
| Nerve Tibial | 47555 | × | - | - | - | - | - | - | - | - |
| Ovary | 4586 | Ovary | 346550 | 69500 | 276899 | × | 155632 | 315170 | 39533 | 99286 |
| Pancreas | 17451 | Pancreas | 252470 | 2004 | 281414 | × | 81234 | 256002 | 51479 | 218849 |
| Pituitary | 6703 | × | - | - | - | - | - | - | - | - |
| Prostate | 4410 | × | - | - | - | - | - | - | - | - |
| Skin Not Sun Exposed Suprapubic | 22215 | × | - | - | - | - | - | - | - | - |
| Skin Sun Exposed Lower leg | 41294 | × | - | - | - | - | - | - | - | - |
| Small Intestine Terminal Ileum | 3500 | Small Intestine | 270628 | 170557 | 171197 | × | 189419 | 139581 | 45684 | 53091 |
| Spleen | 8307 | Spleen | × | 13501 | 315506 | × | 124962 | 267783 | 153962 | 213867 |
| Stomach | 14100 | Stomach Mucosa | × | 111365 | 212099 | 84198 | × | 67217 | 46348 | 17149 |
| Testis | 31732 | × | - | - | - | - | - | - | - | - |
| Thyroid | 47647 | × | - | - | - | - | - | - | - | - |
| Uterus | 2398 | × | - | - | - | - | - | - | - | - |
| Vagina | 2554 | × | - | - | - | - | - | - | - | - |
| Whole Blood | 35143 | × | - | - | - | - | - | - | - | - |

Table S5: Summary of the number of labeled nc-sindels in 15 immune cell types in DICE

| Tissue | #Sample Size |
|---|---|
| Naive B cells | 12312 |
| Naive CD4+ T cells | 14342 |
| Stimulated CD4+ T cells | 9419 |
| Naive CD8+ T cells | 14976 |
| Stimulated CD8+ T cells | 9845 |
| Non-classic monocytes (M2) | 11791 |
| Classic monocytes (monocytes) | 12835 |
| NK cells | 9302 |
| Tfh | 14019 |
| Th1 cells | 10451 |
| Th17 cells | 15317 |
| Th2 cells | 14470 |
| Th1/17 cells (THSTAR) | 13296 |
| Memory Treg cells | 13750 |
| Naive Treg cells | 15496 |

### 2 Supplementary figures

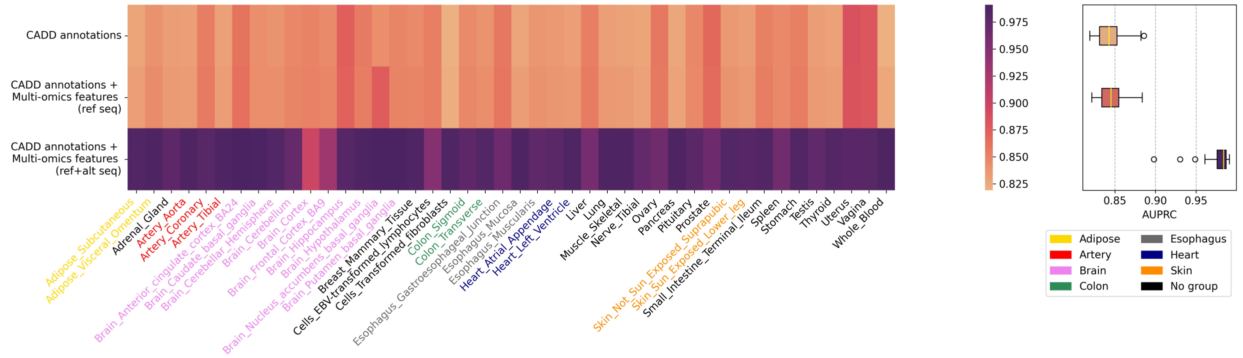


Figure S1: Comparison of three feature sets of TIVAN-indel: (i) 45 CADD functional annotations; (ii) 45 CADD functional annotations+919 multi-omics features under reference sequence; (iii) 45 CADD functional annotations+919 multi-omics features under both reference and alternative sequence across 44 tissues in GTEx. The median of AUPRC across 44 tissues is reported for each feature set.

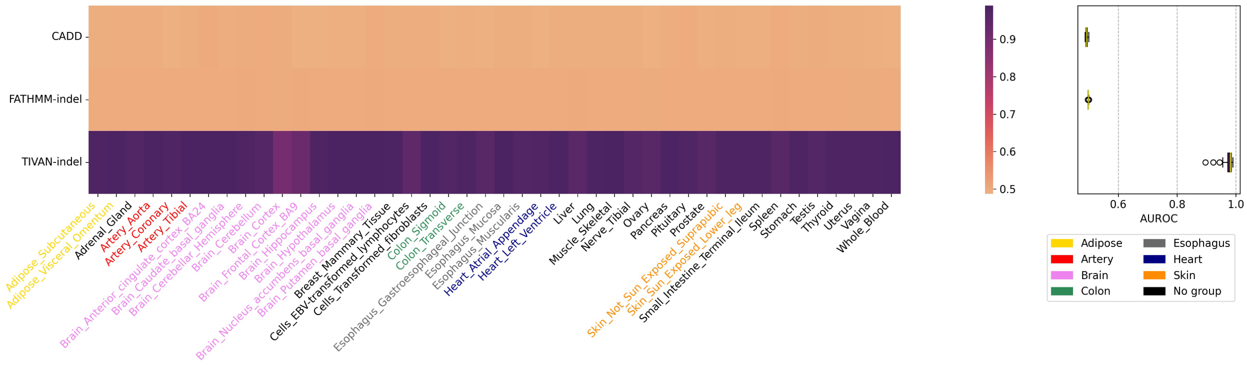


Figure S2: Comparison between TIVAN-indel, CADD and FATHMM-indel for 44 tissues/cell types in GTEx using the precomputed scores from CADD and FATHMM-indel. AUROC is reported for each tissue/cell type.

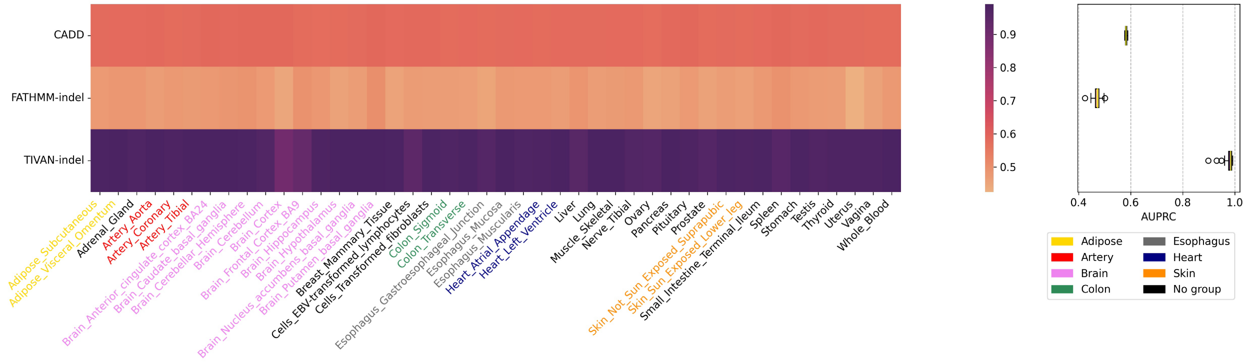


Figure S3: Comparison between TIVAN-indel, CADD and FATHMM-indel for 44 tissues/cell types in GTEx using the precomputed scores from CADD and FATHMM-indel. AUPRC is reported for each tissue/cell type.

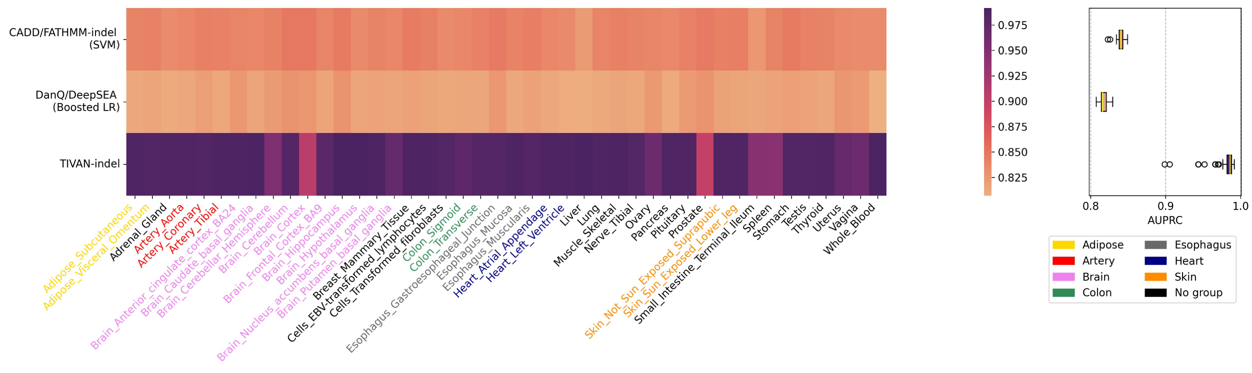


Figure S4: Comparison between TIVAN-indel, CADD/FATHMM-indel (SVM) and DanQ/DeepSEA (Boosted LR) using the within-tissue approach for 44 tissues/cell types in GTEx, where each method is trained and tested using the five-fold cross-validation. AUPRC is reported for each tissue/cell type.

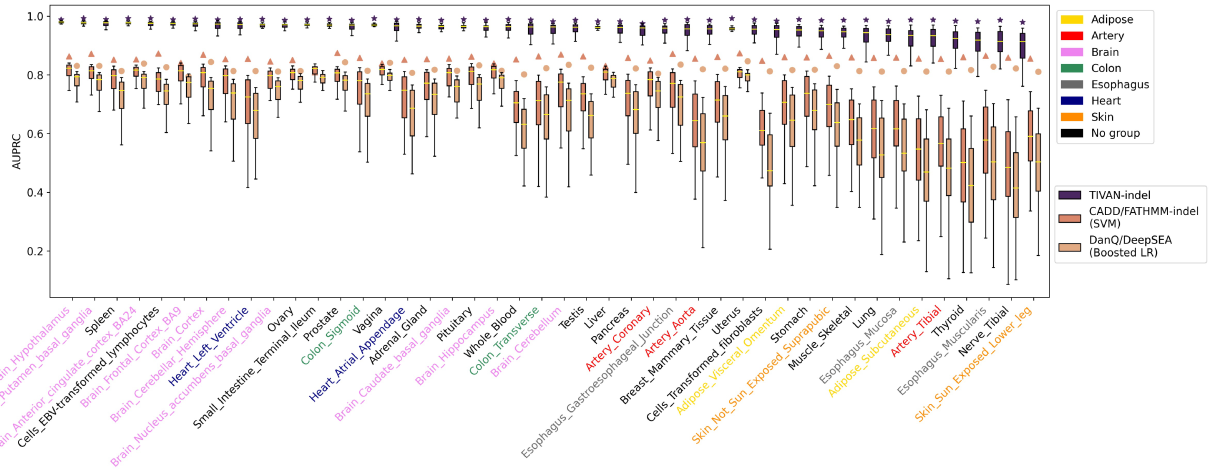


Figure S5: Comparison between TIVAN-indel, CADD/FATHMM-indel (SVM) and DanQ/DeepSEA (Boosted LR) using the independent cross-tissue approach for 44 tissues/cell types in GTEx, where each method is trained using one tissue/cell type and tested on the remaining 43 tissues/cell types. The overlapped samples between the training set and the testing set are removed from the testing set. For each tissue/cell type, AUPRC is reported for the testing 43 tissue/cell types as demonstrated in the boxplot. The asterisk denotes the AUPRC calculated from the cross-validation within-tissue approach. 44 tissues/cell types are colored in 8 tissue/cell types class.

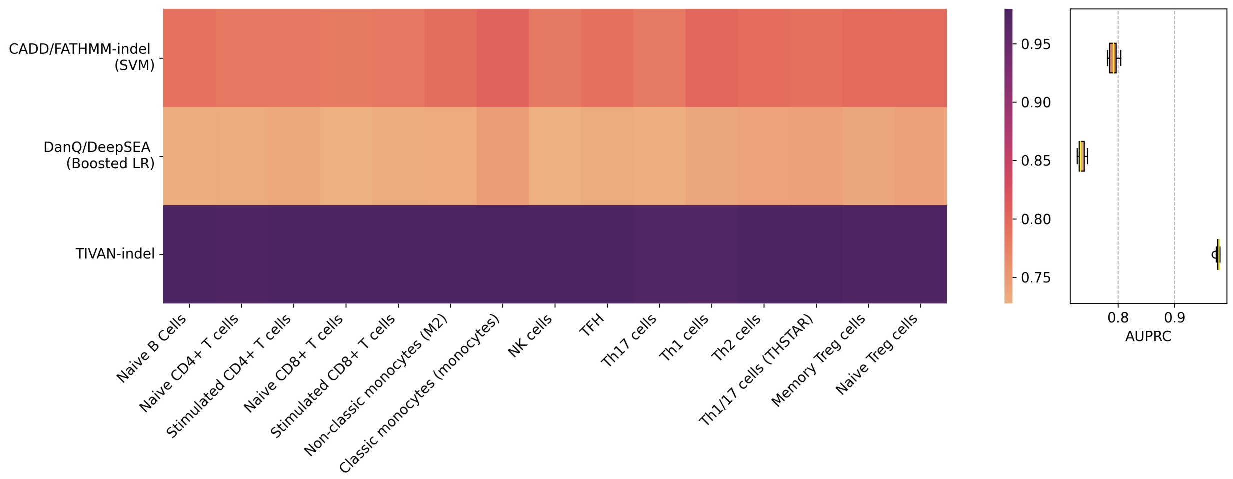


Figure S6: Comparison between TIVAN-indel, CADD/FATHMM-indel (SVM) and DanQ/DeepSEA (Boosted LR) by training on tissue “Whole blood” in GTEx data and testing on the regulatory nc-sindels of 15 immune cell types in DICE. The overlapped samples between training and testing sets are removed from the testing set. The AUPRC is reported for each cell type.

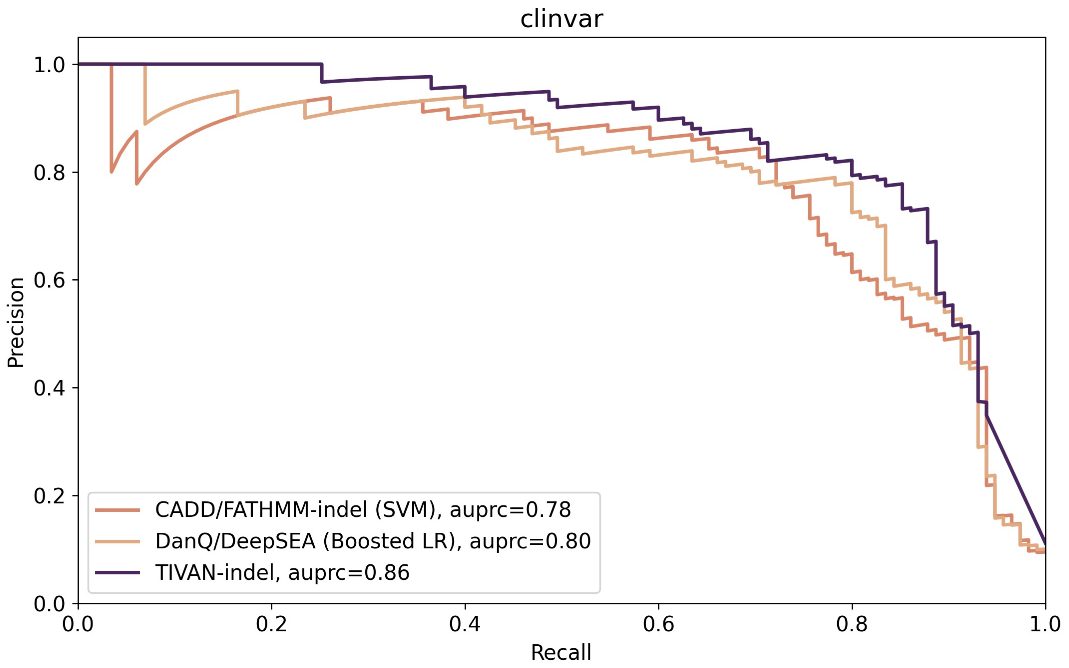


Figure S7: Comparison between TIVAN-indel, CADD/FATHMM-indel (SVM) and DanQ/DeepSEA (Boosted LR) on classifying pathogenic and benign nc-sindels in ClinVar. The AUPRC is reported.
